# Optimised control of adaptive evolution with competing selective pressures

**DOI:** 10.1101/2024.04.08.588561

**Authors:** Marco Corrao, Gabriel Abrahams, Harrison Steel

## Abstract

The development of methods to understand and control the population dynamics of microbial evolution remains an outstanding question in synthetic biology and biotechnology more broadly. Due to the stochastic nature of evolution, its limited observability, and complex intra-population dynamics, this presents a significant challenge. In this paper, we explore techniques to control the evolutionary dynamics of a population, based on manipulation of one or two orthogonal selective pressures, which may in turn be coupled to mutagenesis. Our approach builds on past research in evolutionary biology that developed frameworks to study intra-population variant competition during asexual adaptive processes (i.e. clonal interference). Extending this theory, we design optimal control strategies for one (or more) selective pressures that can be used to maximise the rate of adaptation across a population as a whole. We introduce a theoretical modelling framework for this process, which we support with both simulations and preliminary experimental data, providing a concrete basis for emerging control approaches to directed evolution and evolution-aware design.

## I. INTRODUCTION

Continuous adaptive laboratory evolution (ALE) has become an extensively used tool in modern biotechnology applications [1, 2]. In a typical setup, a microbial population is propagated for prolonged periods of time under controlled conditions. Over the course of the experiment mutant lineages spontaneously emerge and sub-populations with improved fitness are naturally selected for, so that the mean fitness of the population steadily increases over time. By growing the population in a selective environment, the fitness of an individual can be coupled to a phenotype of interest, enabling ALE to optimise a wide range of biotechnologically relevant traits.

In recent years, the fast development of laboratory automation technology has drastically increased the throughput at which evolution experiments can be run, making it possible to run continuous, automated evolution experiments with minimal manual intervention. Particularly, the recent development of programmable continuous growth devices [3, 4] enables a range of experimental parameters to be tightly controlled over the course of an experiment. Therefore, a natural question is how these parameters should be selected and/or controlled to optimise the evolution rate of a population in an ALE experiment.

In this context, a fundamental issue is to understand how the rate of adaptation of an asexual population depends on key parameters of the system, such as the population size, *N*, the *per capita* beneficial mutation rate, *U*_*b*_, and the distribution of fitness effects of acquired mutations. When mutations are rare and population sizes moderate, mutants that have acquired beneficial mutations generally fix in the population far before new mutants can emerge (where we consider a mutation as having *fixed* if it is present in a large fraction of the population and becomes a common ancestor of all future populations). In this Strong-Selection-Weak-Mutation (SSWM) regime the adaptive walk proceeds as a sequence of isolated selective sweeps. Hence, adaptation rate is primarily determined by the influx of new beneficial mutations in the population and, assuming a model where all mutations have the same selection coefficient *s*, the rate of fitness increase is proportional to *NU*_*b*_*s*^2^ [5]. However, in many practical applications, populations are large and/or mutations are sufficiently common such that new mutant lineages typically establish before earlier ones can fix - defined as the Weak-Selection-Strong-Mutation (WSSM) regime. In asexual populations, where recombination is absent, these lineages will compete with each other for fixation – a phenomenon known as *clonal interference*. In this interference regime, travelling-wave models have been a popular framework [5–9] to study the dependence of the adaptation rate on system parameters. Typically, these models approximate the fitness distribution in the population as a stationary wave travelling at constant speed. The dynamics of the bulk of the population are modelled deterministically. whereas the fittest edge of the distribution - the small number of recently arisen mutants that have greater than average fitness - is given detailed stochastic treatment. The speed of the travelling wave is then obtained, implicitely or explicitly, by equating the rates of adaptation implied by these two analyses (for a full treatment of the matter, see [5]).

The dynamics of clonal interference become particularly relevant to practical applications whenever multiple, competing selective pressures operate in the same experiment (Fig. 1). As an example, if *in addition* to a desired selective pressure an experimentalist introduces factors such as UV light or chemical mutagens to increase mutation rates in a population, a second competing pressure would be added to the system. This arises because these mutagenic sources are often toxic to the cell, meaning tolerance-conferring mutations can arise and provide mutants with a selective advantage. If not controlled appropriately, the strength of this secondary selective force can become comparable or even greater than the primary one, effectively reducing the rate and/or trajectory of evolution of the trait of interest in the experiment.

**Fig. 1.**
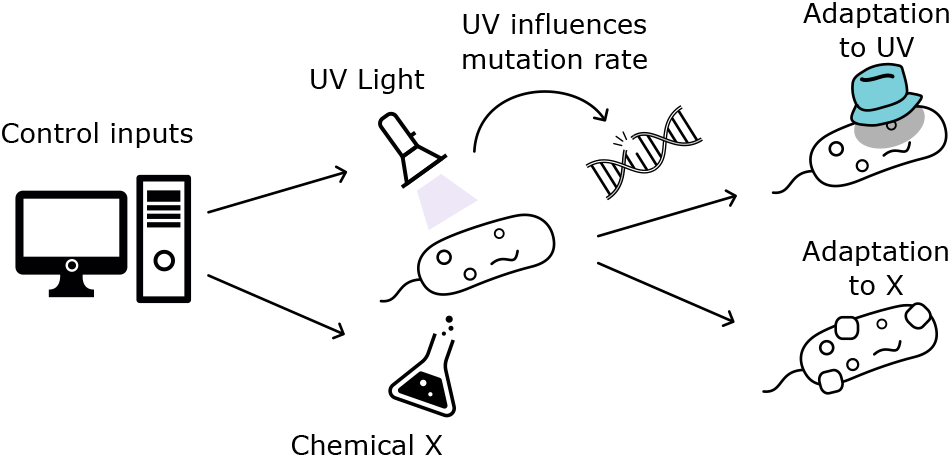
In each generation of the evolutionary process, a mutation establishes that improves adaptation to the selective pressures present (here UV and chemical X). One pressure (here UV) is chosen such that it also influences the mutation rate. The pressures can be controlled to optimally steer the evolutionary trajectory for adaptation of a desired trait (here resistance to chemical X).

In this work, we take initial steps to model and analyse optimal control strategies for such evolutionary processes, in which two coupled mutational processes compete with each other. To this end, we consider a single-effect-size travelling wave model of asexual adaptation [5] and extend it to account for the presence of two competing processes. By computing the probability of fixation for each mutant type, an expression is derived for the adaptation velocity of each process under interference. Hence, treating the selective pressure for each adaptive process as an experimentally controlled input, an optimisation problem is formulated so as to maximise the rate of mutation accumulation in the adaptive direction of primary interest. We derive analytical approximations for this optimum as a function of the system’s parameters, and validate our results against stochastic Fisher-Wright simulations of adaptative dynamics.

## II. Results

### A. Population Model

Our model is developed based on the approach of Desai and Fisher [5]. We first briefly introduce their model, before departing by extending it to consider the interactions between two orthogonal traits. Note that our abbreviated introduction of the modelling framework serves only to provide the necessary context for our work - for a detailed explanation of all terms and relevant derivations (for the single-trait case) please refer to Desai and Fisher [5]. As usual for population models, at small population sizes, the dynamics must be treated stochastically, while at large population sizes, they can be treated deterministically. While calculating the time-dependent probability distributions governing the size of mutant sub-populations is complex, Desai and Fisher [5] showed that by calculating an (approximate) transition time between the stochastic mutant birth-death process, and the subsequent deterministic sub-population growth, it is possible to derive analytical results that agree well with simulation. This approach is particularly useful for our work, as it allows us to find closed-form expressions to guide our intuition regarding how system parameters can be optimised.

Consider a sub-population of *n* individuals of a larger total population of size ∑_*i*_ *n*_*i*_ = *N* (Fig. 2). This sub-population’s fitness, *r*, is defined as the difference between its growth rate and the mean growth rate over the total population *N*. A deterministic sub-population will therefore grow in frequency within the larger population at rate *dn/dt* = *rn* (equivalently a sub-population with below-average fitness −*r* would *reduce* in frequency as *dn/dt* = −|*r*| *n*). Similarly, a stochastic sub-population will follow a simple birth-death process, with birth rate 1 + *r* and death rate 1. Beneficial mutations occur at rate *U*_*β*_ , and increase the fitness of the sub-population from which they arise by *s*. We assume both that all mutations have the same fixed effect size *s*, and there are no deleterious mutations (i.e. *s >* 0). These assumptions are motivated by past work [5, 9]: allowing for variable effect sizes, one finds that the beneficial mutations which tend to dominate the process are narrowly distributed around a size 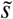, and the impact of deleterious mutations has in *most* cases minimal impact on the overall process dynamics. Defining the most fit sub-population to have fitness *r* = *qs*, Desai and Fisher [5] showed that

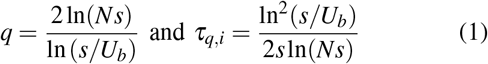

where *τ*_*q*,*i*_ is the establishment time for a population to arise and become deterministic in isolation. This “nose” population (with fitness *qs*) is treated stochastically. It is is fed by a deterministic “feeder” subpopulation with fitness (*q* − 1)*s* (Fig. 2).

**Fig. 2.**
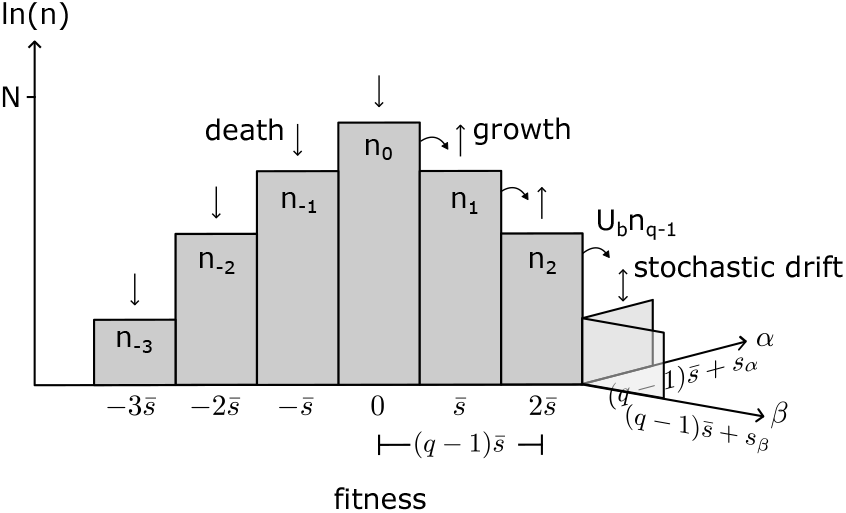
Subpopulations of average size *n*_*i*_ with fitness 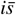 make up the total population of size *N*, with *n*_0_ ≈ *N*. At the nose, we examine how the two possible leading subpopulations feed into the leading background population. Populations with fitness 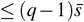 grow deterministically, while the nose populations grow stochastically.

Departing from the above single-trait process, in our model there are two traits being selected for, referred to as type *α* and *β* . Each has a single (but potentially different) fitness effect size *s*_*α*,*β*_ and beneficial mutation rate *U*_*α*,*β*_ . Considering mutations of type *α* and *β* arising from a common background (quantified below), we wish to calculate the probability *x* that the type *β* of these two mutants becomes fixed, conditional on the assumption that one of these two fixes in the population as a whole. To proceed we consider the “race” to fixation in two parts. First is establishment, a period of length *τ*_*q*_ during which population dynamics are stochastic, and will typically be “won” by the mutant type with greater beneficial mutation rate *U* . Following establishment is a period of approximately deterministic growth, favouring whichever mutant provides the larger fitness benefit (i.e. greater *s*_*α*,*β*_ ). The rate at which fixed mutations are acquired (i.e. combining both processes) is given by 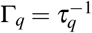, and the rate at which *β* mutations are acquired is given by

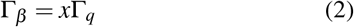

which we later seek to optimize.

Deterministic growth of each mutant sub-population begins following their establishment, at which they have reached size 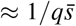 where we define 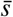 as the mean effect size of mutations that fix in the population i.e.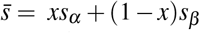 . Following establishment each has a growth advantage of 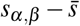compared to the mean mutation effect size 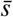 for mutants arising from the common background 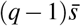 (see Fig. 2). This growth advantage is compounded over the time taken to go from initial establishment to the point at which each population reaches its maximum size (i.e. fixation, where population size is on the order of ≈ *N*): this takes (*q* − 1) establishment times, hence (*q* −1)*τ*_*q*_ in total. After this period mutant *α* will have a size advantage of 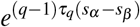 compared to type *β* . If a mutant of type *β* (rather than one of type *α*) is to fix it therefore needs to have a similarly sized head-start at the time of establishment. Thus, we approximate mutant *β* as having “fixed” if the following condition is satisfied:

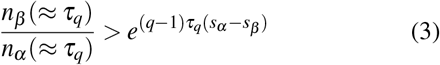

. Conceptually (3) is stipulating that if (for example) *s*_*β*_ *< s*_*α*_ , then a mutation of type *β* needs to reach establishment quickly and grow by a factor of 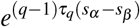 before a type *α* mutant establishes, lest it be overtaken in the race to fixation (due to the *α* mutant’s faster deterministic growth).

We now calculate the probability of the condition in (3) being satisfied following establishment. As we are primarily analysing the (stochastic) dynamics of establishment, we approximate both mutant types as having the same fitness during establishment (i.e.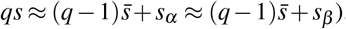, but allow (potentially) different mutation rates. By making this assumption we will observe that calculation of (3) becomes independent of time.

Following from (3) we express the probability of a *β* mutation achieving the requisite “head start” as 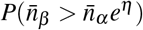 where *η* = (*q* − 1)*τ*_*q*_(*s*_*α*_ − *s*_*β*_ ) and 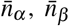 are independent random variables. Assuming that the probability of *β* fixing is less than 50% (i.e. 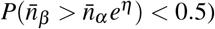 we arrive at the following expression:

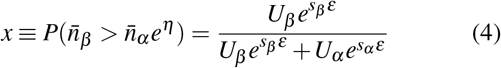

where *ε* = (*q* − 1)*τ*_*q*_.

### B. Control Parameters

So far, we have assumed that the effect sizes of new mutations, *s*_*α*_ and *s*_*β*_ , are fixed constants. In practice, however, the selective advantage of a mutation is dependent on the growth environment and, as such, can be tuned by the experimentalist. For example, if the primary evolution process under study involves increasing the tolerance to an inhibitory compound, selective pressure in the system can be modulated by varying the concentration of this compound in the growth medium [10, 11]. In order to model this possibility, we introduce two nondimentional parameters 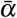 and 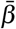, with 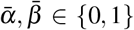, corresponding to the level of selective pressure applied on each adaptation process. To simplify our analysis, we shall assume that selection acts linearly on the system, so that the fitness effect size for each process becomes

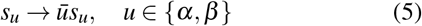

Conceptually parameters 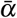 and 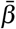 are defining an *environment*, and imply that mutations arising (which we are approximating as having an *identity* independent of the environment) nevertheless have a different *effect* (in terms of their fitness benefit) depending on these parameters.

At the same time, increasing the selective pressure will cause a proportionate, overall slow-down in growth, which we model by rescaling the adaptation rates as

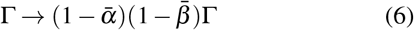

To gain an understanding of the above model, it’s useful to consider some limiting cases, using the *β* mutation process as an example. At very low selective pressure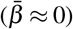, the population grows approximately at its maximal rate, but mutations introduce negligible fitness gains as there is no pressure to which they can beneficially adapt, and so adaptation is virtually absent. Conversely, at nearly maximal selective pressure 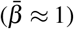, fitness gains from new mutations approach their maximum, but the population grows vanishingly slow, so that the timescale for the establishment and fixation of new mutants become overwhelmingly large. Intuitively, then, fitness-effect sizes and growth slow-down trade-off with each other – if growth is inhibited there is conversely a large room- for-improvement due to mutations – suggesting the existence of an optimal level of selective pressure in between these limits.

Finally, we consider the case where the two processes under investigation are mutationally coupled. Practically, this happens if a toxic mutagenic source, such as UV light, is applied to the system, so that one of the two processes (which we here assign to the *α* direction) involves evolution of tolerance to this input. Under these circumstances, increasing 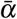 (which corresponds, for example, to increasing the intensity of UV irradiation on the system) will also have the effect of increasing global mutation rates. Assuming again a linear effect, we thus have:

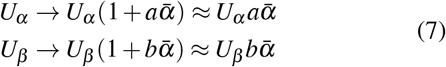

where *a*,*b* are proportionality constants and the second approximation in (7) follows from assuming that we operate in a regime where induced mutagenesis contributes the majority of mutations to each process.

### C. Maximising Adaptation Rate

By introducing coupling between the two processes, the mutagenic input 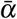 affects evolution in the *β* direction both positively, by increasing the influx of *β* -type mutations, and negatively for *two distinct reasons*: it introduces a competing selective advantage for mutations in the *α* direction, and *also* slows growth overall according to (6). Assuming *β* to be the process of primary experimental interest, we consider the problem of finding an optimal input, 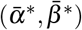, such that:

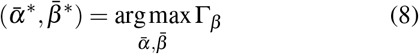

In order to derive an explicit expression for *v*_*β*_ in terms of the control inputs, we note that, under the optimal operating conditions, adaptation rate in the *α* direction should typically be smaller than in the *β* direction, so that, in this regime, the combined rate term in (2) can be approximated by the rate in the *β* direction when present in isolation:

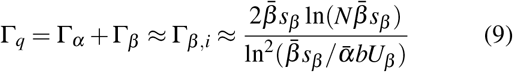

where we obtained the last equality substituting (5) and (7) into (1). By the same argument, we can expect that at the optimum,

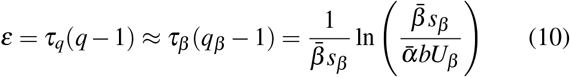

where the last equality follows from (1). Substituting these two approximations into (2), and using the transformations (5) and (7), we obtain an explicit, approximate expression for Γ_*β*_ valid in the vicinity of the optimum:

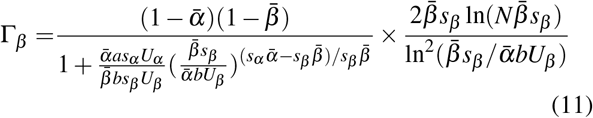

Introducing the lumped parameters

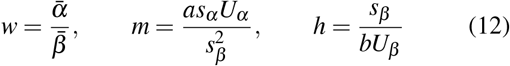

Equation (11) can be rewritten in the form

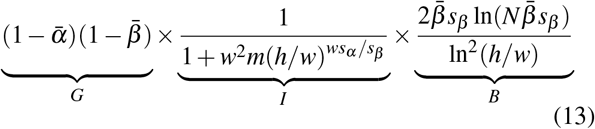

Here, we have split up the expression for Γ_*β*_ into three conceptual parts. The *G* term represents growth slow-down due to the application of each pressure. The sigmoidal *I* term models velocity slow-down arising from *α* type mutations interfering with *β* type during the fixation process. Finally, the *B* term corresponds to the adaption velocity the *β* process would have if operating in isolation (but with selection coefficient and mutation rate scaled by the choice of control parameters), thus providing an upper bound on the magnitude of Γ_*β*_ .

With this expression at hand, we now proceed to study the behaviour of the optimal input pair 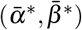 as a function of the remaining system’s parameters. We start by focusing our attention on the primary selective pressure 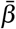. In the vicinity of the optimum, we expect interference from the *α* process to be always rather contained, so that the dominant contributions to the expression in (13) are given by the *G* and *B* terms. In addition, for the typical parameter range considered here (where *N* ≫ *s*_*β*_ and *s*_*β*_ */U*_*b*_ ≫ 1), we have that *Ns*_*β*_≫ 1 and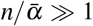, so that the two logarithms in the *B* term depend only weakly on 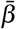. As a result, we find that near the optimum the velocity for the primary process obeys

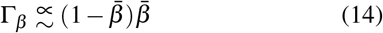

which is maximised for

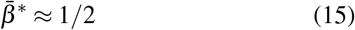

. Hence, to the level of approximation considered here, the optimal input for the primary process can be treated as independent of the system’s parameters. The approximate independence of the optimal 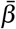 was verified via stochastic Fisher-Wright type simulations of the model (Fig. 3).

**Fig. 3.**
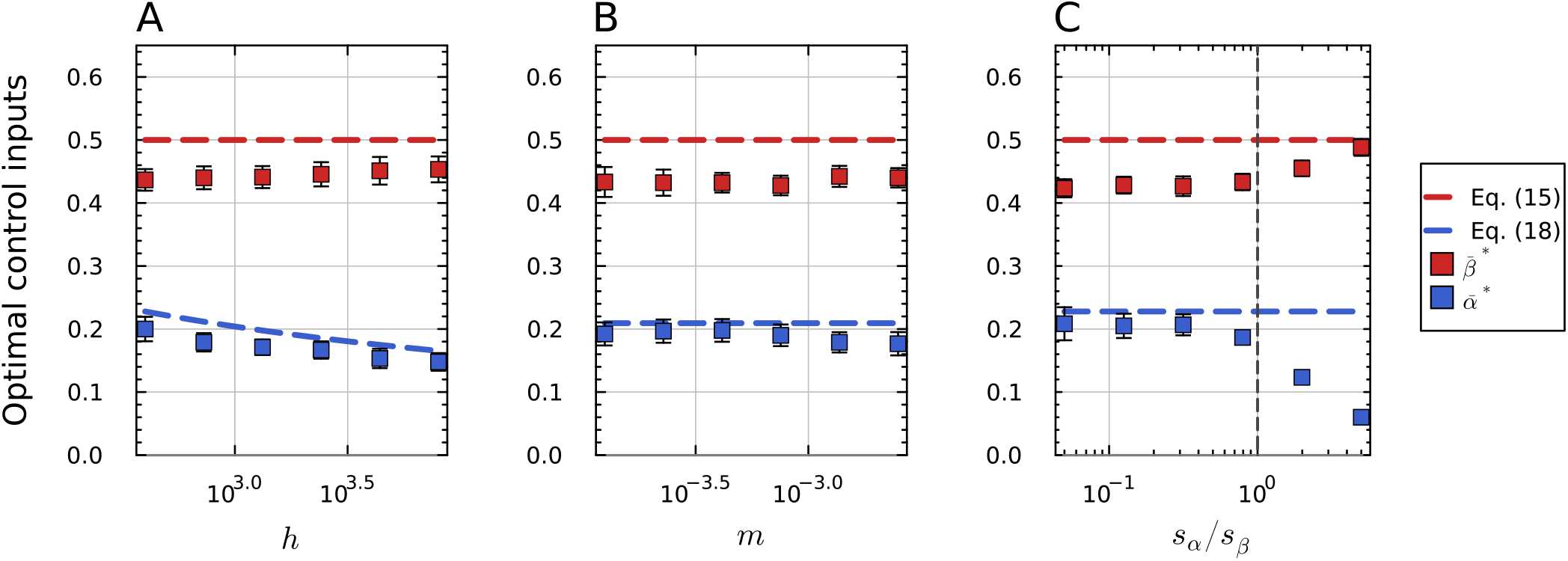
Optimal control policies as a function of model parameters. Squares represent numerical estimates computed using a grid search over a discretisation of the 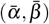 space. For each choice of policy, the adaptation rate in the *β* direction was computed by averaging the output of 50 stochastic Fisher-Wright simulations of the population’s dynamics, using an adapted simulation algorithm from [9]. Markers show the mean and standard deviation of 15 independent replicates of this grid search. Dashed lines indicate the analytical approximations provided in the main text. A: Optimal policy as a function of *h* = *s*_*β*_ */*(*bU*_*β*_ ), with *s*_*α*_ = *s*_*β*_ = 0.02, *U*_*α*_ = 5 × 10^−7^, and *U*_*β*_ varied between 10^−6^ and 5 × 10^−8^. B: Optimal policy as a function of 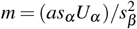, with *s*_*α*_ = *s*_*β*_ = 0.02, *U*_*β*_ = 5 10^−7^, and *U*_*α*_ varied between 10^−6^ and 5 10^−8^. C: Optimal policy as a function of *s*_*α*_ */s*_*β*_ , with *U*_*α*,*β*_ = 10^−6^, *s*_*β*_ = 0.02, and *s*_*α*_ varied between 0.001 and 0.1. The vertical black line denotes the limit beyond which our analytical approximations are no longer valid. For all three cases, we set *N* = 10^10^ and *a* = *b* = 50.

Next, we consider the input for the mutagenic process,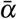 . Noting that, with the exception of the *G* term, expression (13) depends on 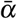 only through the ratio of selective pressures *w*, we turn our attention to the equivalent problem of finding the optimal value for this parameter. This will give us an indication of how strong the mutagenic pressure should be set relative to the pressure of the desirable adaptation process *β* . Differentiating (13) with respect to *w* and maintaining only leading order terms, we get that the optimal *w* approximately solves:

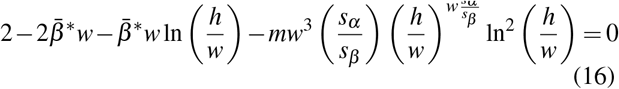

When *s*_*α*_ ≤ *s*_*β*_ (i.e. when the nominal selection coefficient for the mutagenic process is smaller or comparable in magnitude to the one for the main process), the above equation is dominated by the first two terms near the optimum, which we expect to find in the region where *w <* 1. Neglecting this contribution, we find that the solution to the resulting transcendental equation can be given in terms of the Lambert-W function, with the optimal ratio *w*^*^ being approximately:

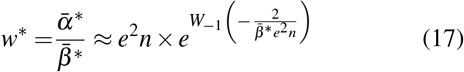

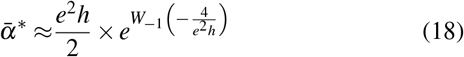

where *W*_−1_(*z*) denotes the −1 branch of the Lambert W-function the second approximation follows by substituting in our previous result 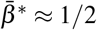. Hence, in this regime, the optimal input of the mutagenic process depends solely on the properties of the main adaptation process, showing a weak, negative dependence on 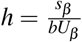 (Fig. 3). For *s*_*α*_ *> s*_*β*_ , the interference term in equation (16) is no longer negligible even at the optimum, and the above approximation ceases to hold. In this parameter regime, numerical experiments show a fast decrease of the optimal mutagenic input profile, which intuitively is required to dominate the increasing competitiveness of *α*-type mutations in the race to fixation with the *β* -type ones.

Following the above analysis we have derived (approximate) expressions for optimal values of 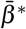 and 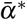 which if achieved would (in principle) maximally accelerate theaccumulation of mutations adapting the population to stressor 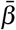; this is illustrated by simulations for varying control parameters in Fig. 4.

**Fig. 4.**
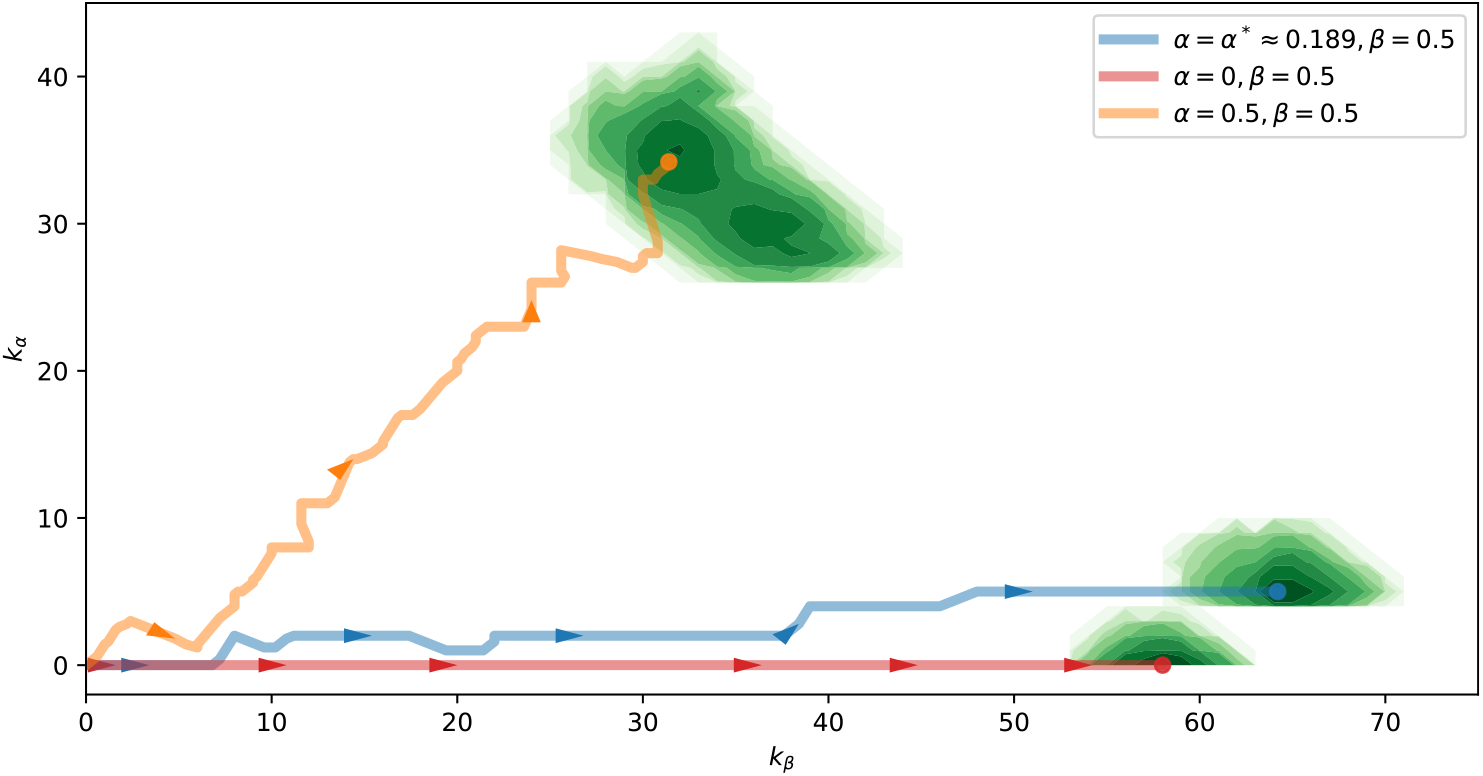
Simulated evolution trajectories under the optimal (blue) and various suboptimal (orange, red) control policies. Here, the continuous lines track the change over time in the mean number of each mutation type in the population. The full distribution of mutation frequencies within each population at the final time point is shown via the green density plots for completeness. Each trajectory is of the same length in time, as measured by the number of generations scaled by the slow-down factor 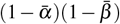 . Simulation parameters are *N* = 10^9^, *U*_*α*,*β*_ = 10^−5^ and *s*_*α*,*β*_ = 0.01. The optimal trajectory (blue) fixes a larger number of *β* type mutations (compared to the 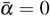 case, red) due to 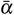 increasing the value of *U*_*β*_ , *despite* it also fixing ≈ 5 *α* type mutations. However, for excessively high 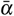 (orange) the population fixes a large number of *α* type mutations, which reduces accumulation in the *β* direction due to clonal interference.

## III. DISCUSSION

The ever-increasing availability and accessibility of laboratory automation has opened a range of opportunities for improvements in the field of laboratory evolution. Among these, the automated control of selective pressure has been experimentally validated as a powerful way of accelerating adaptation in continuously-cultured microbial populations [10, 11]. In these experiments, an exponentially growing population is kept at near-constant size by continuous dilution and its growth rate measured throughout. Using an appropriately designed feedback controller, the growth environment is then dynamically modulated to maintain growth at a constant fraction of its uninhibited value, resulting in a sustained selective pressure driving the adaptation process. In this paper, we took initial steps to develop the theory and control approach necessary to derive an optimal value for this inhibition setpoint by considering a simple model of mutation-fixation dynamics in the the presence of tunable selective strength. We generalised our discussion by considering the scenario where multiple, coupled adaptive processes operate in the same experiment.

If only one adaptive process is present (which corresponds to the limiting case of *U*_*α*_ = 0 or 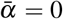 in our analysis), we find that the model admits an optimal growth inhibition setpoint (corresponding to 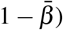 ) which, for the parameter range investigated here, remains approximately constant at half of the uninhibited growth rate of the populations. To begin exploring this finding in an experimental setting, we adapted three *E. coli* populations to UV stress in the Chi.Bio bioreactor platform [4], each with a differently tuned PI controller trying to maintain its growth rate at ≈ 1*/*3 (i.e. just below half) of its base-line growth rate (that observed in the absence of any selective pressure). This is shown in Fig. 5. We found that adaption proceeded faster (as measured by the time taken for the population to tolerate the maximum deliverable UV input level) as the controller was tuned to better maintain the setpoint, suggesting promising future validation experiments in this direction.

**Fig. 5.**
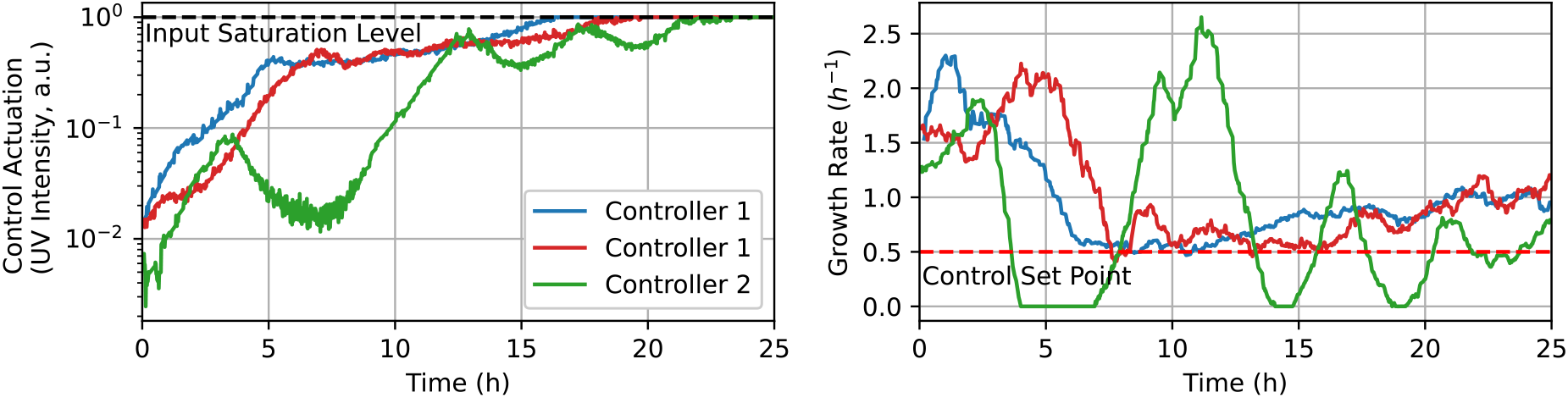
Adaptation to UV stress for three independent *E. coli* populations in the Chi.Bio bioreactor platform. In each experiment, the UV irradiation intensity is modulated by a PI controller (each with different tuning) with the goal of maintaining the growth of the culture at the desired setpoint. To account for the fact that adaptation of microbial populations typically proceeds multiplicatively with respect to the absolute stressor level, the output of the controller was exponentiated. Left: UV (285nm) input provided by each controller, measured relative to the maximum deliverable irradiation power. Right: measured growth rate from each reactor, showing different variable performance among controllers in maintaining the setpoint. The control set-point is set to 1/3 of the average observed initial growth-rate (≈ 1.5 *h*^−1^)

When a mutagenic process with toxicity is added to our system (i.e. input 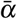 ), we find that gains in the adaptation rate of the primary process can be obtained by increasing the mutagenic input, up to an optimal value. Past this optimum, interference from mutants tolerant to the mutagenic source becomes dominant and adaptation in the primary direction rapidly slows down. For this process, the optimum level depends on the parameters of the model. However, for a range of parameter values of experimental interest, we found this dependence to be rather weak, suggesting that, in practice, an appropriate value can be found even if the parameters of the evolution process are not known or measurable.

Intuitively, the diminishing returns observed through increasing 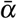 to boost *U*_*β*_ are expected - the benefit of increasing 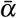 is limited due to the weak logarithmic dependence on mutation rate (i.e. in Equation (1)) when populations are in a regime where mutations are abundant (i.e. *NU*_*b*_ ≫ 1), as this regime is characterised by processes that are *already* being saturated due to competition between different co-existing lineages. Future work will investigate other regimes of this process; for example, the case in which process *β* is initially in the Strong-Selection-Weak-Mutation regime characterised by the more forgiving (with regard to mutation rate dependence) adaptive velocity *NU*_*b*_*s*^2^. Further, by extending these results to the travelling-wave frameworks in which mutation sizes (*s*) are drawn from a continuous distribution [9] it will be possible to determine optimal control policies that additionally bias a mutation-selection process toward fixation of mutations of specific size range/s.

## Notes

* All authors contributed equally to this work. M.C. and G.A. acknowledge support from the BBSRC Interdisciplinary Bioscience DTP [grant number BB/T008784/1]. H.S. is supported in part by EPSRC projects EP/W000326/1 and EP/X017982/1.

### Competing Interest Statement

The authors have declared no competing interest.

